# Systematic identification of human SNPs affecting regulatory element activity

**DOI:** 10.1101/460402

**Authors:** Joris van Arensbergen, Ludo Pagie, Vincent FitzPatrick, Marcel de Haas, Marijke Baltissen, Federico Comoglio, Robin van der Weide, Hans Teunissen, Urmo Võsa, Lude Franke, Elzo de Wit, Michiel Vermeulen, Harmen Bussemaker, Bas van Steensel

**Author notes:** Correspondence should be addressed to J.v.A. or B.v.S.

## Abstract

Most of the millions of single-nucleotide polymorphisms (SNPs) in the human genome are non-coding, and many overlap with putative regulatory elements. Genome-wide association studies have linked many of these SNPs to human traits or to gene expression levels, but rarely with sufficient resolution to identify the causal SNPs. Functional screens based on reporter assays have previously been of insufficient throughput to test the vast space of SNPs for possible effects on enhancer and promoter activity. Here, we have leveraged the throughput of the SuRE reporter technology to survey a total of 5.9 million SNPs, including 57% of the known common SNPs. We identified more than 30 thousand SNPs that alter the activity of putative regulatory elements, often in a cell-type specific manner. These data indicate that a large proportion of human non-coding SNPs may affect gene regulation. Integration of these SuRE data with genome-wide association studies may help pinpoint SNPs that underlie human traits.

## INTRODUCTION

About 100 million single-nucleotide polymorphisms (SNPs) have been identified in human genomes \ The vast majority of these are located in non-coding regions, and in a typical human genome about 500 thousand SNPs overlap with candidate regulatory elements such as enhancers and promoters ^1^. It is becoming increasingly clear that such non-coding SNPs can have substantial impact on gene regulation ^2^, thereby contributing to phenotypic diversity and a wide range of human disorders ^3–5^.

Genome-wide association studies (GWAS) are a powerful tool to identify candidate SNPs that may drive a particular trait or disorder ^6,7^ Similarly, expression quantitative trait locus (eQTL) mapping can identify SNPs that are associated with the expression level of individual genes ^3,8^. Unfortunately, even the largest GWAS and eQTL studies rarely achieve single-SNP resolution, largely due to linkage disequilibrium (LD). In practice, tens to hundreds of linked SNPs are correlated with a trait. Although new fine-mapping techniques ^9,10^, integration with epigenome data ^11^, deep learning computational techniques ^12^ and GWAS studies of extremely large populations can help to achieve higher resolution, pinpointing of the causal SNPs remains a major challenge.

Having a list of all SNPs in the human genome that have the potential to alter gene regulation would mitigate this problem. Ideally, the regulatory impact of SNPs would be measured directly, but due to the vast number of human SNPs, this requires high-throughput assays. Two methods have been employed for this purpose. First, changes in chromatin features such as DNase sensitivity and various histone modifications have been mapped in lymphoblasts or primary blood cells derived from sets of human individuals with fully sequenced genomes ^13–19^. Here, the chromatin marks serve as proxies to infer effects on regulatory elements, with the caveat that a change in regulatory activity may not always be detected as a change in chromatin state, or vice versa. Furthermore, many traits do not manifest in blood cells, and other cell types are more difficult to obtain for epigenome mapping.

An alternative and often-used functional readout is to insert DNA sequence elements carrying each variant into a reporter plasmid. Upon transfection of these plasmids into cells, the promoter or enhancer activity of these elements can be measured quantitatively. Different cell types may be used as models for corresponding tissues in vivo. Large-scale versions of this approach are referred to as Massively Parallel Reporter Assays (MPRAs). So far, MPRAs have been successfully applied to screen tens of thousands of SNPs ^20–24^. Each of these studies has yielded tens to at most several hundreds of SNPs that significantly alter promoter or enhancer activity. As these MPRA studies have covered only a tiny fraction of the genome, it is likely that many more SNPs with regulatory impact are to be discovered.

So far, the scale required to systematically cover the millions of human SNPs has not been achieved. Here, we report application of an MPRA strategy with a >100-fold increased scale compared to previous efforts. This enabled us to survey the regulatory effects of nearly 6 million SNPs in two different cell types, providing a resource that helps to identify causal SNPs among candidates generated by eQTL and GWAS studies.

## RESULTS

### A survey of 5.9 million SNPs using SuRE

We applied our Survey of Regulatory Elements (SuRE) technology to systematically screen millions of human SNPs for potential effects on regulatory activity. SuRE is a MPRA with sufficient throughput to query entire human genomes at high resolution and high coverage ^25^. Briefly, random genomic DNA (gDNA) fragments of a few hundred base pairs are cloned into a promoter-less reporter plasmid that, upon transfection into cultured cells, only produces a transcript if the inserted gDNA fragment carries a functional transcription start site (TSS) (Figure 1a). The transcript is identified and quantified by means of a random barcode sequence that is unique for every insert, allowing for a multiplexed readout of millions of random DNA fragments. Importantly, because active promoters as well as enhancers generate transcripts, activity of both types of elements can be assayed quantitatively by SuRE ^25^.

**Figure 1.**
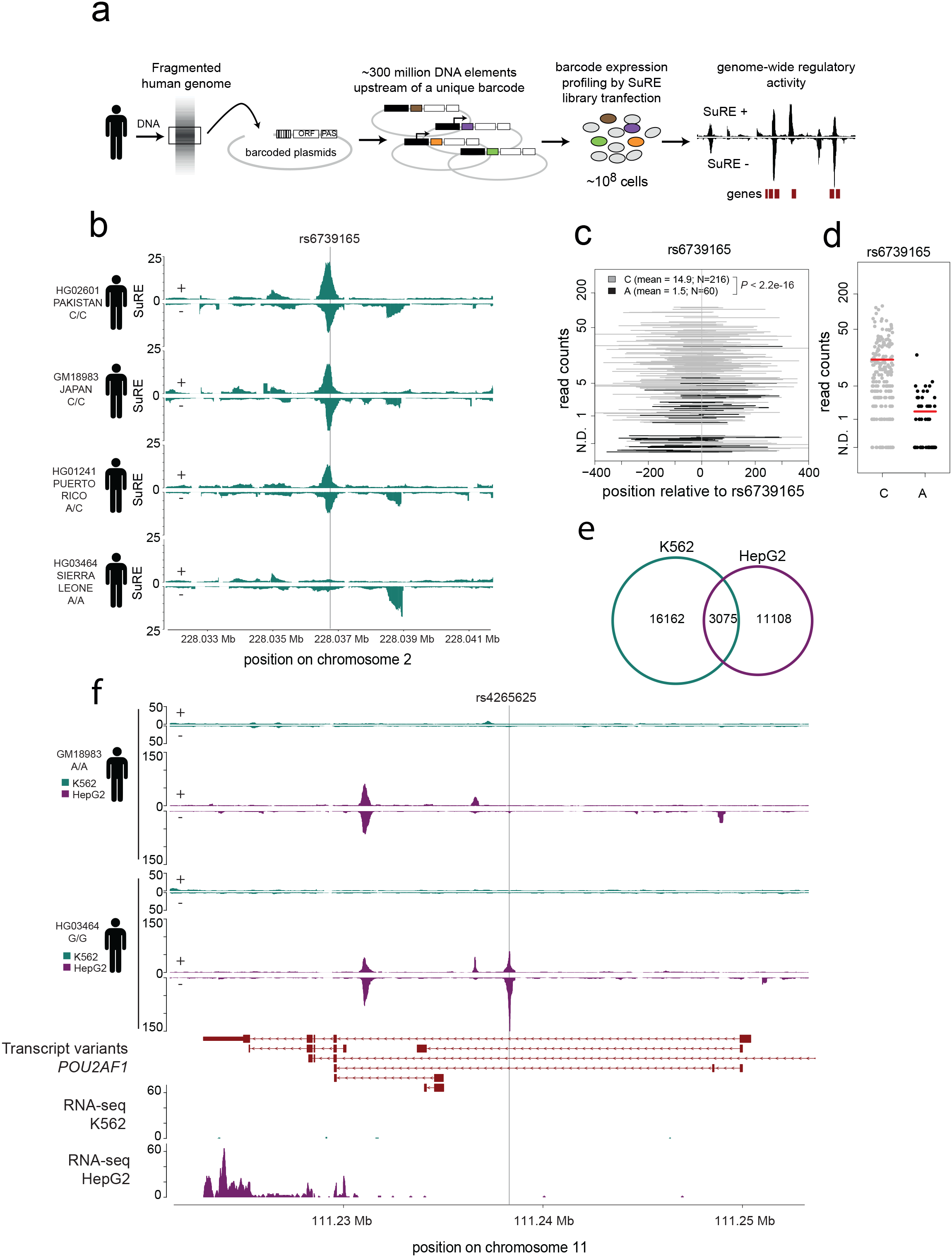
Identification of raQTLs by SuRE. **a**. Schematic representation of the SuRE experimental strategy. ORF, open reading frame; PAS, polyadenylation signal. Colors indicate different barcodes. **b**. SuRE signals from the four genomes in an example locus, showing differential SuRE activity at raQTL rs6739165, depending on the allele (A or C) present. **c**. SuRE activity for all fragments containing rs6739165. **N**.D., not detected. Values on the y-axis were shifted by a random value between -0.5 and 0.5 in order to better visualize DNA fragments with the same value. **d**. Same data as in (**c**), but only the expression value for each fragment. Red lines indicate mean values. **e**. Numbers of raQTLs in K562, HepG2, or both. **f**. Example of a locus showing differential SuRE activity for 2 genomes in HepG2 only. Below the SuRE tracks known transcript variants of *POU2AF1* are indicated, and RNA-seq data from K562 and HepG2 (data from ^27^).

To survey a large cross-section of SNPs present in the human population, we chose four divergent genomes that were fully sequenced by the 1000 Genomes Project ^1^ (Figure 1b). From each genome we generated two independent SuRE libraries that each contained ~300 million random gDNA fragments of 150-500 bp (Figure S1a; **Supplemental Table 1**). In these libraries a total of 2,390,729,347 unique gDNA fragments were sequenced from both ends, mapped to the reference genome and linked to their unique barcode. Among these fragments, 1,103,381,066 carried at least one SNP for which we identified both alleles in our libraries. These libraries enabled us to test promoter/enhancer activity of both alleles of 5,919,293 SNPs, which include 4,569,323 (57%) of the ~8 million known common SNPs (minor allele frequency [MAF] > 5%) world-wide ^1^. Importantly, each variant is covered by 122 different gDNA fragments on average (Figure S1b,c), which provides substantial statistical power.

We introduced these libraries by transient transfection into human K562 and HepG2 cells, which both are well-characterized cell lines that can be transfected with high efficiency. K562 is an erythroleukemia cell line with strong similarities to erythroid progenitor cells ^22^. HepG2 cells are derived from a hepatocellular carcinoma, and are an approximate representation of liver cells. After transfection of the SuRE libraries into each cell line we isolated mRNA and counted the transcribed barcodes by Illumina sequencing. Three independent biological replicates yielded a total of 2,377,150,709 expressed barcode reads from K562 cells and two biological replicates yielded 1,174,138,611 expressed barcode reads from HepG2 cells.

### Identification of thousands of SNPs with regulatory impact

From these data we first constructed tracks of SuRE enrichment profiles for each of the four genomes (Figure 1b). This revealed thousands of peaks that generally colocalize with known enhancers and promoters (Figure S1d), as reported previously ^25^. For a subset of peaks, the magnitude varied between the four genomes and showed a correlation with a particular allele of a coinciding SNP. For example, in K562 cells we detected a strong SuRE signal overlapping with SNP rs6739165 (located in the *COL4A3* gene) that is homozygous for the C allele, but no signal in the genome that is homozygous for the A allele, and an intermediate signal in the genome that is heterozygous for this SNP (Figure 1b).

In order to systematically annotate SNPs we combined for each transfected cell line the complete SuRE datasets from the four genomes. For each SNP we grouped the overlapping gDNA fragments by the two alleles, as illustrated in Figure 1c,d. This allowed us to identify SNPs for which fragments carrying one variant produced significantly different SuRE signals compared to those carrying the other variant. Because all of these fragments differ in their start and end coordinate, the activity of each variant is tested in a multitude of local sequence contexts, providing not only statistical power but also biological robustness. For each SNP we calculated a P-value by a Wilcoxon rank sum test, and compared it to P-values obtained by a random permutation strategy to estimate false discovery rates (FDR) (Figure S1e,f). We also calculated the average SuRE signal (i.e. enrichment over background) for each allele and focused on SNPs for which the strongest allele showed a signal of at least 4-fold over background. We refer to the resulting SNPs at FDR < 5% as *reporter assay QTLs* (raQTLs).

This analysis yielded a total of 19,237 raQTLs in K562 cells and 14,183 in HepG2 cells (Figure 1e). The average allelic fold change of these SNPs was 4.0-fold (K562) and 7.8-fold (HepG2) (Figure S1g,h). In 72% of cases the SuRE effect could be assigned to a single SNP; when SNPs were spaced less than ~200bp apart, their effects could typically not be resolved (Figure S1i).

Most raQTLs were detected in either K562 or HepG2 cells, but not in both (Figure 1e). The overlap may be underestimated due to the somewhat arbitrary FDR and expression cutoffs used to define these sets (Figure S1j). Nevertheless, many SNPs show clear cell type specific effects; for example, rs4265625:G creates regulatory activity in HepG2 only (Figure 1f). Interestingly, rs4265625 lies in *POU2AF1*, a gene that has been linked to primary biliary cirrhosis – a liver disease – in a GWAS study ^26^.

### raQTLs are enriched for known regulatory elements

We systematically analyzed the overlap of the raQTLs with known regulatory elements in K562 cells ^27^. Compared to randomly sampled SNPs, raQTLs showed 5-15 fold enrichment for promoter and enhancer related chromatin types, and depletion for repressed or transcribed chromatin types (Figure 2a). We also observed strong enrichment of raQTLs in DHSs (Figure2b,c), which is consistent with their overlap with enhancers or promoters.

**Figure 2.**
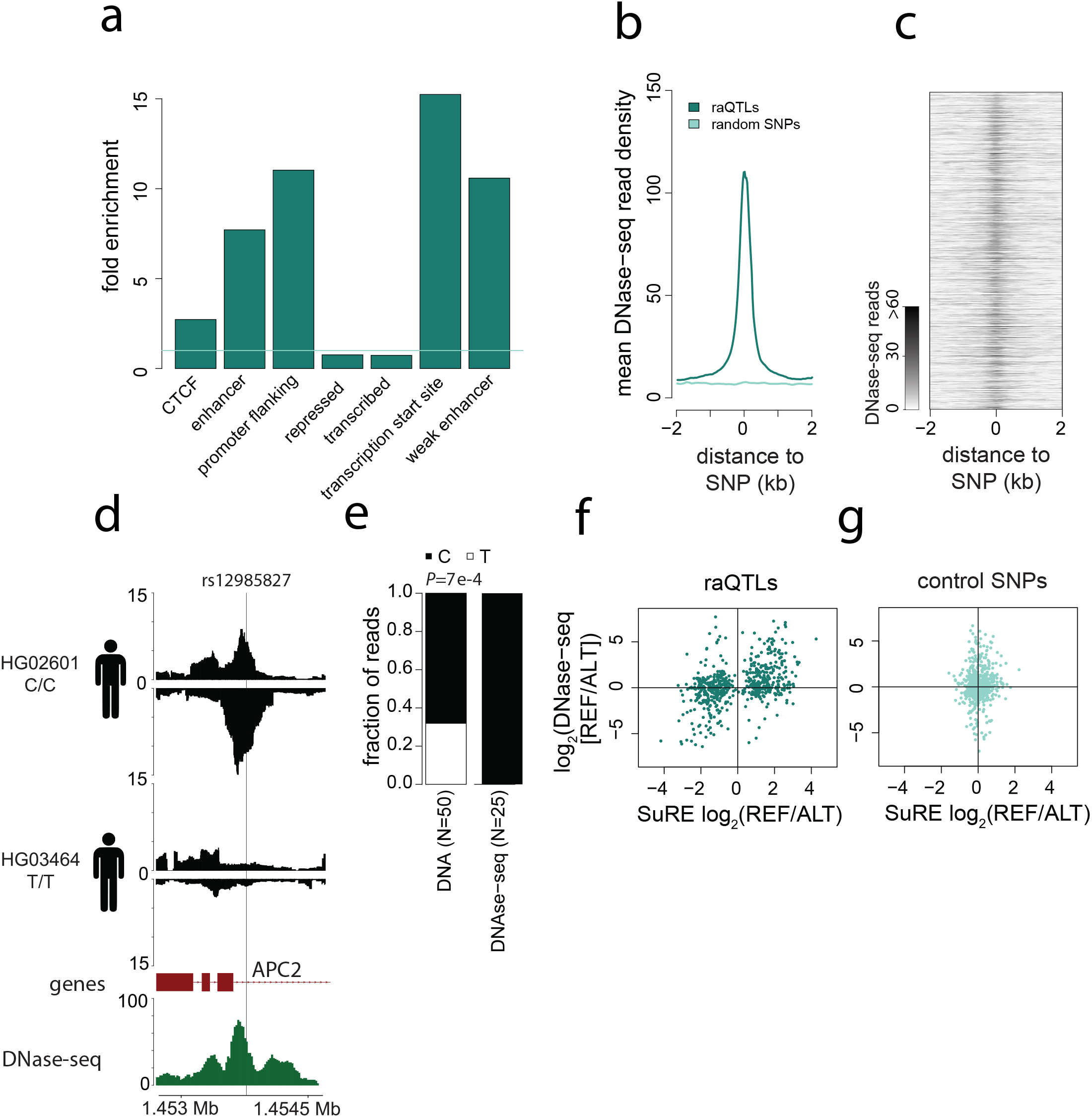
Correlation of SuRE signals with local chromatin states. **a**. Enrichment or depletion of raQTLs among major types of chromatin in K562 ^27^ relative to random expectation. All values are significantly different from 1 (P < 2.2 e-16, Fisher exact test). **b**. Average profile of DNase-seq enrichment for the 19,237 raQTLs (dark green) compared to an equally sized random set of analyzed SNPs (light green). **c**. DNase-seq signals aligned to the 19,237 raQTLs, sorted by their P-value (lowest P-value on top). **d-e**. Example of a SNP with differential SuRE activity for the two alleles, overlapping with a DNase-seq peak in K562 cells (**d**) and showing only DNase sensitivity for one allele, even though both alleles are present in K562 cells (**e**). **f**. Comparison of allelic imbalance of SuRE signals and DNase-seq signals (normalized for genomic DNA allele counts) for 616 raQTLs for which K562 cells are heterozygous. REF: reference allele; ALT: alternative allele. **g**. Same as in (**f**) but for a random set of control SNPs overlapping with a DNase-seq peak. DNase-seq data in b-g are from ^27^.

Some of the raQTLs are heterozygous in the genome of K562 cells. For these SNPs we investigated whether the allelic imbalance observed by SuRE was reflected in a corresponding imbalance in the DHS signal. For example, the SuRE signal at rs12985827, a non-coding variant in an intron of the *APC2* gene, has a strong bias for the C allele over the T allele (Figure 2d), and indeed it shows a similar allelic imbalance for DHS (Figure 2e). Among the 616 raQTLs that were heterozygous in K562 and showed sufficient DNase-seq coverage, we observed a strong skew for higher DNase sensitivity at the more active allele, compared to 616 heterozygous non-raQTL control SNPs that overlapped DHSs (Figure 2f,g). We conclude that available epigenome data is generally consistent with the SuRE results.

### Altered transcription factor binding sites at raQTLs

The observed changes in enhancer or promoter activity are likely explained by SNPs changing transcription factor (TF) binding motifs. For example, the T allele of rs12985827 disrupts an EGR1 binding motif and leads to reduced SuRE activity (Figure 3a, Figure 2d). To investigate this systematically, we made use of the SNP2TFBS database ^28^ that lists computationally predicted changes in TF motifs for most SNPs identified in the 1000 Genomes Project. Among the set of raQTLs as identified by SuRE in K562 and HepG2 cells, 31% and 38% are predicted to alter the motif of at least one TF, respectively (Figure 3b). This is a 1.6-fold and 1.9-fold larger proportion than for all SNPs, respectively. Moreover, for 67% and 69% of the raQTLs (in K562 and HepG2 cells, respectively) there was concordance between the predicted effect on motif affinity and SuRE expression, i.e., the variant with the weakest motif resemblance showed the lowest SuRE expression (Figure 3c). We note that 100% concordance should not be expected in this analysis, for example because not all TF binding motifs are known, and some TFs can act as repressors.

**Figure 3.**
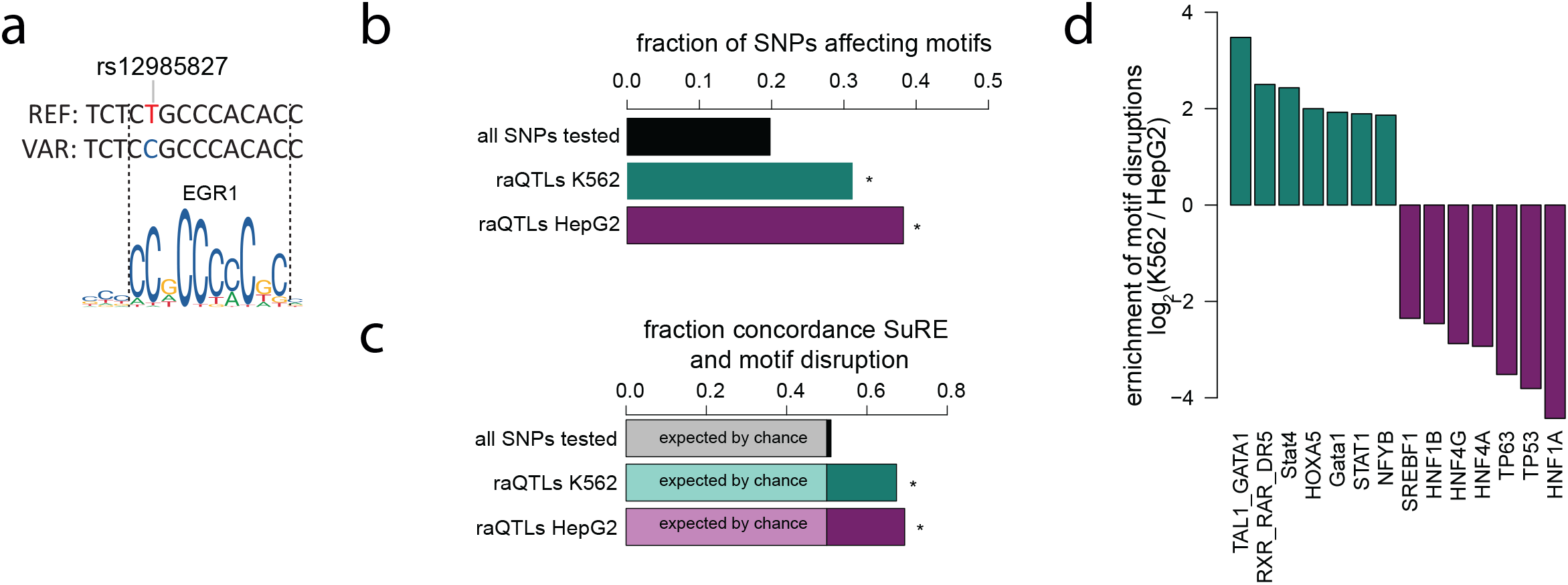
Concordance of SuRE data and predictions based on TF binding motifs. **a**. Comparison of the sequence flanking raQTL rs12985827 (same SNP as in Figure 2d-e) and the sequence logo for EGR1. The T allele disrupts a conserved nucleotide in the EGR1 binding motif. **b**. Compared to all SNPs, raQTLs in K562 and HepG2 both overlap preferentially with computationally predicted alterations of TF binding motifs according to SNP2TFBS ^28^. Asterisks, *P* < 2.2e-16, according to Fisher exact test. **c**. Concordance between the predicted increase or decrease in TF binding according to SNP2TFBS, and the observed effect in SuRE, assuming that decreased TF binding typically leads to decreased activity of a regulatory element. Asterisks, *P* < 2.2e-16, according to Fisher exact test. **d**. TF motif alterations that are preferentially present among raQTLs in either K562 or HepG2 cells. Only the 7 most enriched TF motifs for each cell type are shown.

We expected that motifs in raQTLs reflect the sets of TFs that are selectively active in the respective cell types. Indeed, raQTLs in K562 were enriched for motifs of TFs that are primarily active in the erythroid blood lineage, such as GATA and STAT factors, while raQTLs in HepG2 cells were enriched for motifs of TFs that are specific for liver cells, such as HNF factors (Figure 3d). We also found disruptions of the TP53 motif to be relatively more consequential in HepG2, which is presumably related to the known inactivation of TP53 in K562 cells ^29^ but not in HepG2 cells ^30^.

Together, these data point to a general concordance between the detected changes in SuRE activity and predictions based on sequence motif analysis. We note, however, that computational prediction of TF binding and the downstream functional consequences is still very difficult, and therefore currently no substitute for testing their effect on regulatory activity.

### No evidence for strong negative selective pressure on raQTLs

It has been observed previously that genes that do not have c/s-eQTLs are more likely to be loss-of-function (LOF) intolerant genes, possibly reflecting selection against variants acting upon such genes ^8,31^. To further investigate this, we analyzed the frequency of raQTLs in the proximity of LOF intolerant genes and LOF tolerant genes. We found that the fraction of SNPs that are raQTLs was significantly but only slightly lower in the proximity of LOF intolerant genes than in the proximity of LOF tolerant genes (Figure S2a,b). However, for a set of control SNPs that were matched to the raQTLs for their SuRE activities and coverage in the SuRE libraries we observed a similar pattern (Figure S2a,b). This suggests that the overall density of active regulatory elements, rather than elements affected by SNPs, is lower near LOF genes. Genome-wide, we observed slightly lower minor allele frequencies (MAFs) for our raQTLs as compared to matched SNPs, but only for the K562 dataset and not for the HepG2 dataset (Figure S2c,d). This modest underrepresentation of raQTLs in the human population is consistent with recent computational predictions ^12^ and may point to a slight negative selection pressure. Taken together, we found no evidence for strong negative selective pressure on raQTLs.

### Integration of SuRE with eQTL maps

Next, we integrated our SuRE data with eQTL mapping data from the GTEx Project ^8^. We compared SuRE data from K562 and HepG2 cells with eQTL data from the most closely related tissues, i.e. whole blood and liver, respectively. We note that strong similarity between the two data types should not be expected, because in the GTEx data each gene with significant associations (eGene) is linked to 101 eQTLs on average, of which only one or a few may be causal. Rather, we regard the SuRE data as a filter to identify the most likely causal SNPs among the large number of eQTL candidates.

For each raQTL, the log_2_(ALT/REF) SuRE signal may be concordant with the eQTL signal (i.e., having the same sign as the slope of the eQTL analysis) or discordant (having the opposite sign). The simplest interpretation of concordance is that the SNP alters a regulatory element that positively regulates the eGene; if the variant reduces the activity of the element then it will also reduce the activity of the eGene. In contrast, discordance may point to more indirect mechanisms, e.g., when a SNP alters the promoter of an antisense transcript that in turn interferes with the sense expression of the eGene; or when a SNP alters a promoter that competes with another promoter for a particular enhancer.

Concordant raQTLs are slightly but significantly enriched compared to discordant raQTLs (K562 vs. whole blood: odds ratio 1.13, P = 9e-4; HepG2 vs. liver: odds ratio 1.36, P = 9e-5, Fisher’s exact test). This suggests that direct effects of SNPs on eGenes are somewhat more prominent than indirect effects. Furthermore, discordant SuRE hits tend to be more distant from the TSS of the corresponding eGenes than concordant SuRE hits (K562 vs. whole blood: P = 0.008; HepG2 vs. liver: P = 0.002, Wilcoxon test; Figure S3a,b), in line with the interpretation that discordant SNPs may act indirectly. Because discordant effects are more difficult to interpret, we further focused on SNPs with concordant effects.

### Candidate causal SNPs in eQTL maps and their putative mechanism

Here, we highlight for several physiologically relevant eGenes the most likely causal SNPs based on our SuRE data, and we provide insights into the potential underlying mechanisms.

A first example is *XPNPEP2*, which encodes an aminopeptidase that affects the risk of angioedema in patients treated with an ACE inhibitor ^32^. The GTEx project has linked the expression of this gene in liver to 33 eQTLs, of which 30 are covered by SuRE (Figure 4a). Of these, a single SNP (rs3788853) located ~2 kb upstream of the TSS stands out by a strong (~5-fold) and concordant effect on SuRE activity in HepG2 cells (Figure 4a). Exactly this SNP has previously been demonstrated to alter the activity of an enhancer that controls *XPNPEP2* ^32^. Our independent identification of this SNP indicates that SuRE can successfully pinpoint a functionally relevant SNP among a set of eQTLs.

**Figure 4.**
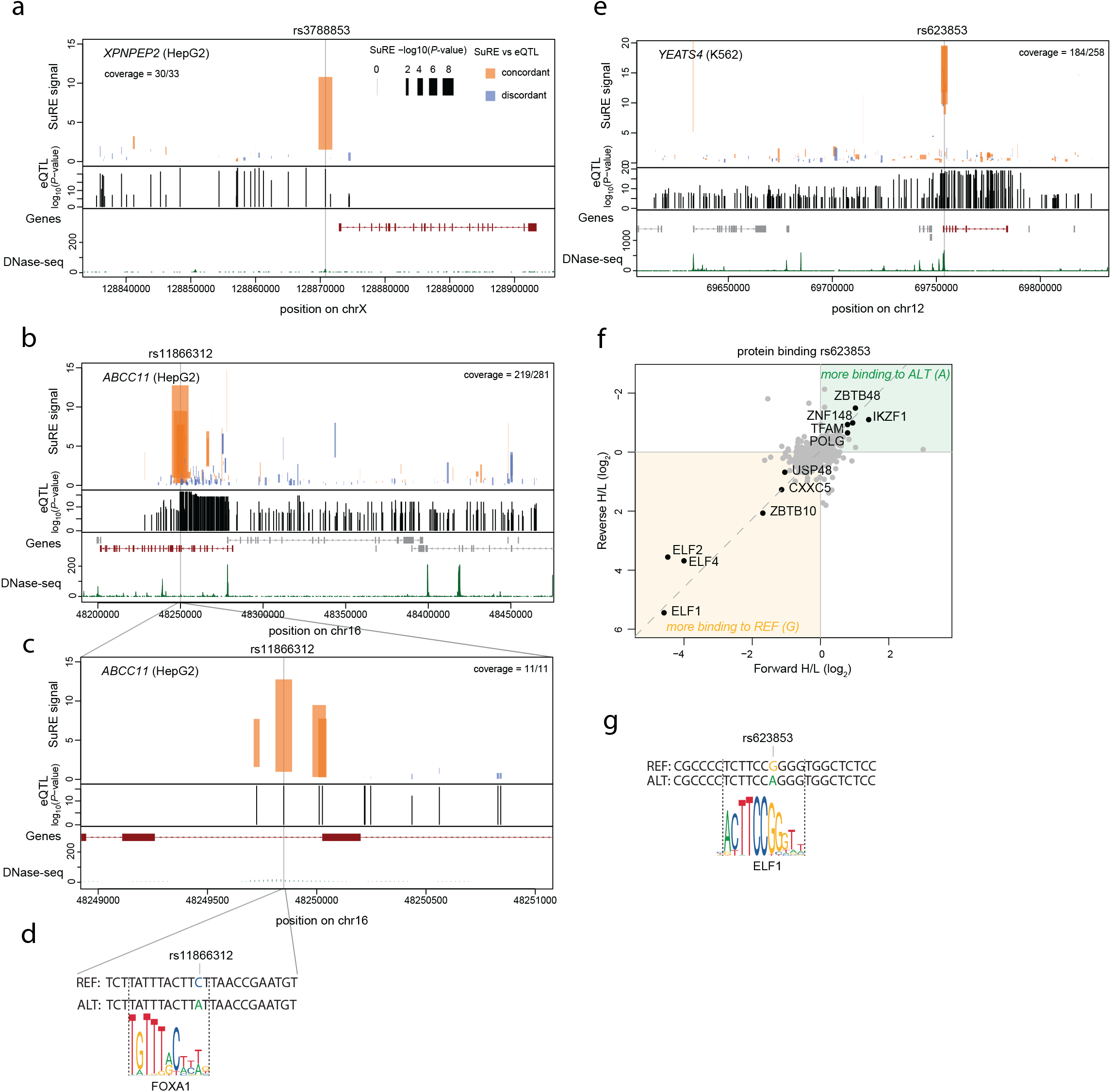
Candidate causal SNPs identified by SuRE among large sets of eQTLs. **a**. Genome track plot showing SuRE signals in HepG2 cells for eQTLs previously identified for the *XPNPEP2* gene in liver according to GTEx v7 ^8^. Top panel: SuRE data in HepG2 cells. The top and bottom of each bar indicate the SuRE signal of the strongest and weakest variant, respectively. Width of the bars is proportional to the -log_10_(P-value); color indicates whether the eQTL effect orientation is concordant (orange) or discordant (blue) with the SuRE effect orientation. Middle panel: positions of significant eQTLs with the associated eQTL -log_10_(P-values). Bottom panel: gene annotation of *XPNPEP2* and DNase-seq data from HepG2 cells ^27^. **b**. Same as (**a**), but for *ABCC11*. **c**. Zoom in of (b). **d**. Sequence of rs11866312 +/- 12 bp aligned to the sequence logo of FOXA1. FOXA1 is predicted to have higher affinity for the A allele than for the C allele. **e**. Same as (**a**) but for *YEATS4* with SuRE data from K562 cells and eQTL data from whole blood (GTEx v7 ^8^). In (**a-c, e**), eGenes are shown in the bottom panels in dark red and all other genes in gray; coverage numbers in the top panels indicate the number of SNPs with SuRE data out of the total number eQTLs. **f**. Mass-spectrometry analysis of proteins from a K562 cell extract binding to 25bp double-stranded DNA oligonucleotides containing either the A or G allele of rs623853. The experiment was performed once with heavy labeling of proteins bound to the A allele and light labeling of proteins bound to the G allele (x-axis), and once with reverse labeling orientation (y-axis). **g**. Sequence of the DNA probes used in (**f**) aligned to the sequence logo of ELF1 ^53^. Consistent with the proteomics analysis, ELF1 is predicted to have higher affinity for the G allele than for the A allele.

A second example is the *ABCC11* gene, which encodes a trans-membrane transporter of bile acids, conjugated steroids, and cyclic nucleotides ^33^. GTEx data from liver identified 281 eQTLs surrounding the *ABCC11* TSS (Figure 4b). SuRE data covered 219 (77.9%) of these, and identified as most likely candidates a cluster of three SNPs within a ~200 bp region located in an intron of *ABCC11* (Figure 4c). Of these, rs11866312 is a likely candidate because the C allele is predicted to disrupt a binding motif for FOXA1 (Figure 4d), a pioneer TF that is highly expressed in liver but not in blood ^8^. Indeed, virtually no SuRE effect of these SNPs is observed in K562 cells (data not shown).

A third example are the neighboring genes *YEATS4* (encoding a transcription regulator) and *LYZ* (encoding lysozyme, an antibacterial protein), which are ~6kb apart. Overlapping sets of eQTLs were identified for these genes in whole blood (Figure 4e and S3c). Among these, SuRE in K562 cells identified two neighboring raQTLs (rs623853 and rs554591) located ~400 bp downstream of the *YEATS4* TSS, which both show concordance with the eQTL data (Figure 4e). To identify transcription factors that might be responsible for the differential SuRE activity, we conducted sequence motif as well as mass spectrometry-based proteomics analyses. For the latter, we incubated biotinylated double-stranded oligonucleotides carrying each of the two variants, immobilized on streptavidin conjugated beads, with nuclear extract from K562 cells. After incubation and washes, we used on-bead trypsin digestion followed by di-methyl stable isotope labeling^34^ and quantitative mass-spectrometry to identify proteins that preferentially associated with one of the two SNP alleles. In this assay, at rs623853 the weaker A allele caused strong loss of binding of Ets-like factors (ELF) (Figure 4f), consistent with a disruption of the cognate motif (Figure 4g). The A allele also showed moderately increased binding of several other proteins. At rs554591 the C allele caused strong loss of ZNF787 and gain of several other factors including KLF and SP proteins (Figure S3d). The variants of both SNPs may thus cause altered binding of TFs and their co-factors, leading to altered enhancer activity. K562 cells are heterozygous for both rs623853 and rs554591, but no significantly different DHS signal is detectable for either of the variants (Figure S3e,f).

These examples illustrate that SuRE can prioritize SNPs as likely causal candidates from a set of tens to hundreds of eQTL candidates. By integrating our data with the GTEx datasets, SuRE identified at least one raQTL among the eQTLs for 20.0% of the 8,661 eGenes in whole blood, and for 11.1% of the 4,000 eGenes in liver. As we illustrated here, sequence motif analysis and mass-spectrometry analysis provide complementary evidence and offer insights into the putative mechanisms.

### Integrating SuRE data with GWAS data

SuRE may also help to identify candidate causal SNPs in GWAS data. To illustrate this, we focused on a large GWAS study that identified more than 1 million SNPs associated with at least one of 36 blood-related traits ^35^. These SNPs occurred in LD-clusters, each on average consisting of 158 SNPs and represented by a single statistically most significant (lead) SNP (i.e. a ‘trait-variant pair’). The lead SNPs are not necessarily the causal SNPs, but are more likely to be close to the causal SNPs. We therefore searched within a 100 kb window from each lead SNP for overlap between significant GWAS SNPs for that trait and significant SuRE hits. For 1,238 out of 6,736 lead SNPs this yielded at least one linked raQTL. Indeed, these raQTLs were preferentially located close to the lead SNPs, compared to a set of matching control SNPs (Figure 5a). Overall, the enrichment of SuRE raQTLs among the total set of GWAS SNPs did not significantly exceed that of the matching control set of SNPs (1188), but this was to be expected considering that only one or a few of the on average 158 significant SNPs may be true causal SNPs.

**Figure 5.**
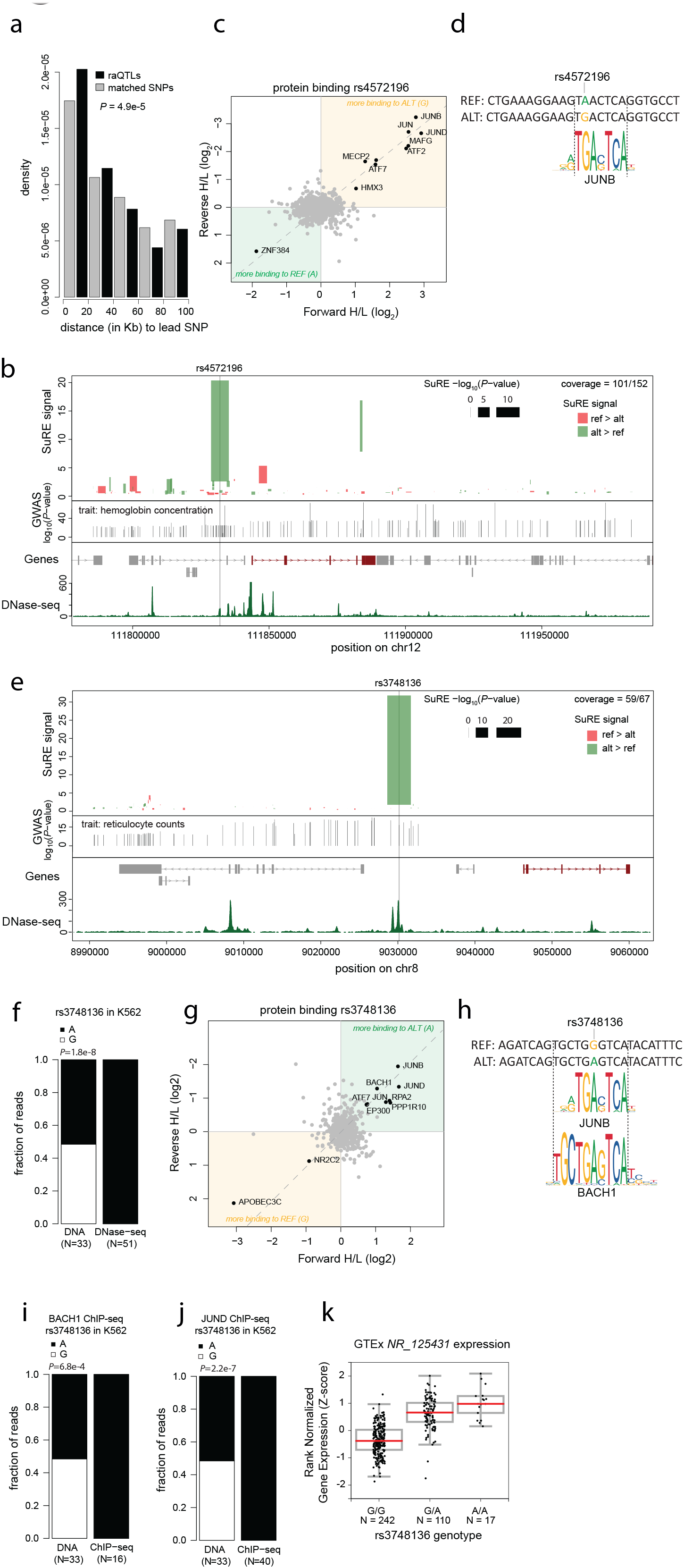
Candidate causal SNPs identified by SuRE among large sets of GWAS SNPs. **a**. raQTLs in K562 cells are modestly enriched in the vicinity of blood GWAS lead SNPs. Plot shows the distribution of distances between lead SNPs for blood traits ^35^ and raQTLs (black) and a set of matched control SNPs (gray). **b**. Genome track plot overlaying SuRE and GWAS data for a cluster of GWAS SNPs linked to hemoglobin concentration ^35^. Top panel: SuRE data in K562 cells. The top and bottom end of each bar indicate the SuRE signal of the strongest and weakest variant, respectively. Color of the bars indicates whether the reference allele (red) or the alternative allele (green) is stronger. Width of the bars is proportional to -log_10_(P-value). Middle panel: positions of significant GWAS SNPs with the associated -log_10_(P-values) ^35^ on the y-axis. Bottom panel: gene annotation track (dark red: *SH2B3;* gray: other genes) and DNase-seq data from K562 cells ^27^. **c**. Protein binding analysis as described in Figure 4f, for rs4572196. **d**. Sequence of the probes used in (**c**) aligned to the sequence logo for JUNB. Consistent with the proteomics analysis, JUNB is predicted to have a higher affinity for the G allele. **e**. Same as (**b**) but for a cluster of SNPs associated with reticulocyte counts by GWAS ^35^. Long non-coding RNA gene *NR_125431* is shown in dark red. **f**. Fraction of reads containing each of the two variants for rs3748136 in K562 genomic DNA and K562 DNase-seq reads. **g**. Same as (**c**) but for rs3748136. **h**. Sequence of the probes used in (**g**) aligned to sequence logos for JUNB and BACH1. Consistent with the proteomics analysis, JUNB and BACH1 are predicted to show more affinity for the ALT allele. **i**. Barplots indicating fraction of reads containing each of the two variants for rs3748136 in K562 genomic DNA (left) and K562 ChIP-seq reads for BACH1 (right). **j**. Same as (i) but for ChIP-seq reads for JUND. ChIP data are from ^27^. **k**. Association between variants of rs3748136 and *NR_125431* expression in whole blood according to GTEx ^8^.

One example where SuRE provided a clear candidate among the GWAS SNPs is rs4572196, which is within 100 kb of 11 lead SNPs that were preferentially associated to various mature red blood cell traits, such as ‘Hemoglobin concentration’, ‘Red blood cell count’ and ‘Hematocrit’ ^35^. Interestingly, rs4572196 lies ~11kb upstream of *SH2B3* (also known as *LNK)*, a gene that regulates megakaryopoiesis and erythropoiesis ^36^. In none of the 11 GWAS associations rs4572196 is the lead SNP, but in SuRE the G allele shows an ~8-fold higher activity than the A allele (P = 2.0e-08) (Figure 5b). K562 are homozygous for the A allele and the region shows a weak signal in DNase-seq (Figure 5b). By *in vitro* proteomics we identified several proteins with differential binding activity to the two SNP alleles. JUN proteins showed about 5-fold stronger binding to the G allele (Figure 5c), which fits with the observation that this G nucleotide is an essential part of the consensus JUN binding motif (Figure 5d). The GWAS study demonstrated a positive association between the reference A allele and hemoglobin concentration and red blood cell counts. SuRE identifies the A allele as the weak allele, potentially reducing SH2B3 levels, which would be consistent with the fact that SH2B3 in mouse inhibits erythropoiesis and thrombopoiesis ^36^. Collectively, these data strongly point to rs4572196 as a candidate causal SNP linked to red blood cell traits, potentially as the result of altered JUN binding.

Another example is rs3748136, which, together with 66 other SNPs in this locus, was found to be linked to blood counts of reticulocytes. Amongst the 59 SNPs covered in our data, rs3748136 is the only significant SuRE hit and has a very large effect size, with the A allele showing an approximately 18fold higher activity than the G allele (p = 7.5e-20) (Figure 5e). K562 are heterozygous for this variant and indeed show a strong allelic imbalance in DHS-seq with the A allele being the more active variant (Figure 5f). To explore the possible mechanism driving this imbalance, we performed *in vitro* binding proteomics analysis on the two alleles. Among others, this identified JUN and BACH1 proteins as more strongly bound to the A allele (Figure 5g), consistent with the G allele disrupting predicted binding motifs for both proteins (Figure 5h). Both BACH1 and JUN proteins are highly expressed in whole blood and in K562 cells ^8,27^. Reanalysis of available ChIP data for BACH1 ^27^ showed a complete allelic imbalance for BACH1 binding and the same was found for JUND (Figure 5i,j). ChIP-seq data for JUNB is currently not available in K562 cells, but JUND binds essentially the same motif. SNP rs3748136 is located in an intergenic region, and eQTL analysis of whole blood ^8^ has linked the A allele to elevated expression of the nearby non-coding RNA gene *NR_125431* (Figure 5k). To further test this, we attempted to modify the G allele in K562 cells to an A allele by CRISPR/Cas9 editing ^37^. These experiments were confounded by apparent genetic instability and substantial clone-to-clone variation in expression of *NR_125431* in K562 cells, both in non-edited and G->A edited cells (Figure S4a-c). However, we did observe an approximate 2.5 fold upregulation for the clones in which we had substituted the G allele for the A allele, with borderline statistical significance (P = 0.03, one-sided Student’s t-test; Figure S4d).

Finally, to illustrate the utility of the HepG2 dataset we overlaid our data with a recent GWAS study on hepatocellular carcinoma (HCC) ^38^. In this work the authors identified variants in the HLA class 1 region that are associated to hepatitis B virus (HBV) related HCC, a type of cancer prevalent in Asia. Within the HLA class 1 region, 61 SNPs were identified in a ~200kb continuous region and 50 of these were present in our dataset, 2 being hits in HepG2 (Figure 6) (and none were hits in K562). Both of these raQTLs are intergenic with the alternative allele being associated with higher HCC risk and lower SuRE signal, suggesting that the alternative alleles disrupt enhancers in the region that play a role the protection against hepatitis B virus related HCC. Besides the HLA genes that are important for the presentation of self and non-self peptides to T cell receptors (TCRs), several other genes in this region could play a role in HCC. For example, ZNRD1 and its antisense transcript have been implicated in expression of HBV mRNA and proliferation of HBV infected HepG2 cells ^39^.

**Figure 6.**
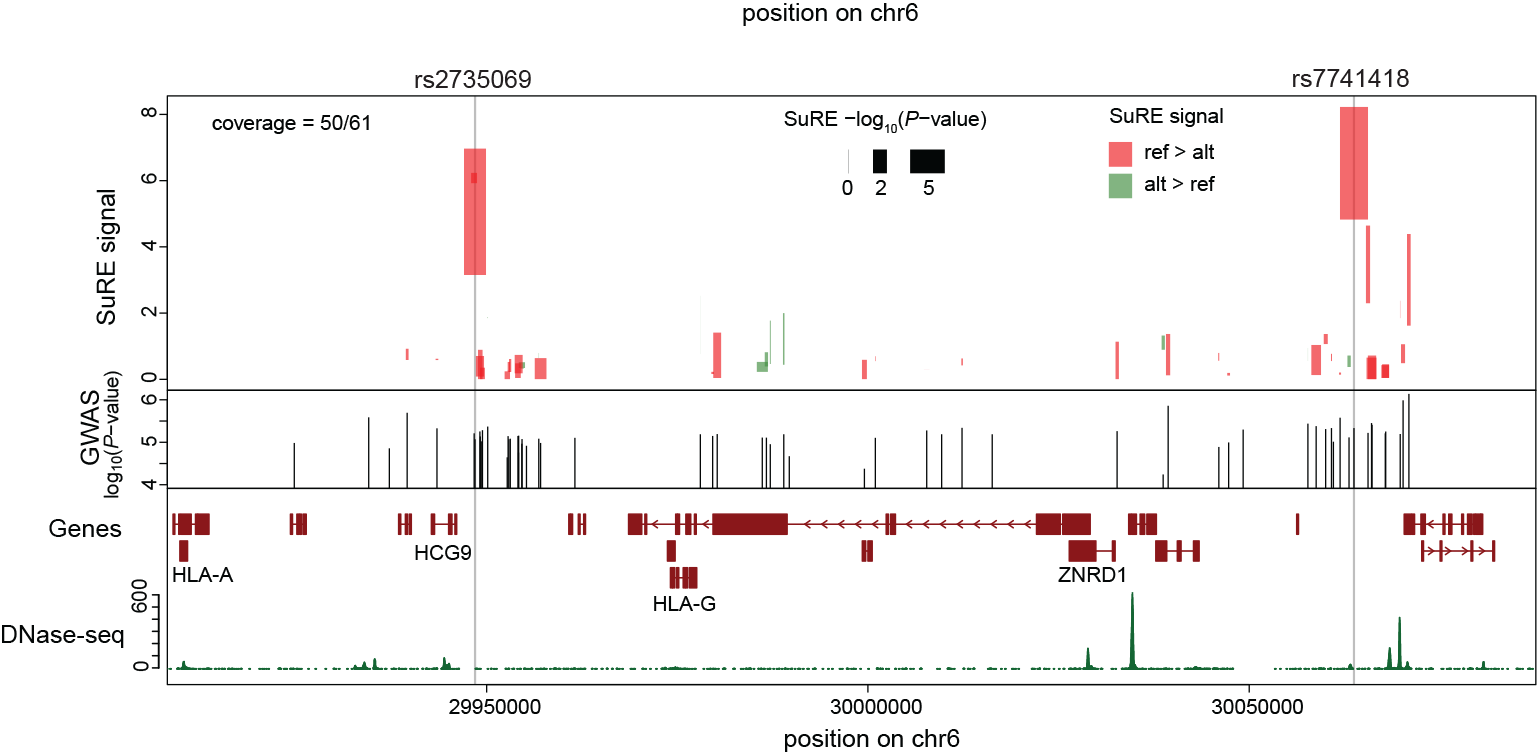
Candidate causal SNPs identified by SuRE among GWAS SNPs for hepatocellular carcinoma. Comparison of SuRE and GWAS data for a cluster of GWAS SNPs linked to Hepatocellular carcinoma ^38^. Top panel: SuRE data in HepG2 cells. The top and bottom end of each bar indicate the SuRE signal of th strongest and weakest variant, respectively. Color of the bars indicates whether the reference allele (red) or the alternative allele (green) is stronger. Width of the bars is proportional to -log_10_(P-value). Middle panel: positions of significant GWAS SNPs with the associated -log_10_(P-values) ^38^ on the y-axis. Bottom panel: gene annotation track and DNase-seq data from HepG2 cells ^27^.

## DISCUSSION

MPRAs have previously been used successfully to generate initial lists of SNPs that can alter the activity of regulatory elements ^20–24^. These earlier efforts, however, covered only a tiny proportion of the total collection of human SNPs. Here, we applied SuRE to increase the scale by >100-fold. By surveying nearly 6 million SNPs from 4 entire human genomes we identified about 30 thousand SNPs that alter regulatory activity of enhancers or promoters. Because 90% of these raQTLs were identified in only one of the two tested cell lines, it is likely that extension of this survey to other cell types will increase the number of raQTLs substantially. It is thus conceivable that several percent of all human SNPs may have impact on the activity of regulatory elements in at least one cell type.

For the purpose of SNP characterization, SuRE differs from previous MPRAs not only in scale, but also in the redundant design. SuRE tests each SNP in a large number of DNA fragments that differ randomly in their start and end position. This increases the odds that a robust and biologically representative measure of SNP effects is obtained.

Nevertheless, like most other MPRAs, SuRE assays all DNA elements in a plasmid context and in cultured cell lines, which may yield different results compared to a proper genomic context and tissue context. As we illustrated here, integration with multiple orthogonal datasets can help provide confidence in the relevance of candidate SNPs. Sequence motif analysis and *in vitro* binding mass-spectrometry can serve this purpose, and in addition provide key insights into the mechanisms by which individual SNPs may act on gene regulation. Importantly, we illustrated that SuRE can help to narrow down sets of candidate SNPs obtained from methods such as eQTL and GWAS that often suffer from limited mapping resolution.

We foresee several additional applications of these SuRE data. First, there are many other eQTL and GWAS studies that may be overlaid with the SuRE maps. Second, in addition to SNPs, small insertions and deletions (indels) may be analyzed. While in human genomes such indels occur at a ~20 fold lower frequency than SNPs ^1^, their individual regulatory impact may be more potent, as they tend to disrupt TF binding motifs more dramatically. Third, our datasets may be useful for studying the regulatory grammar of TFs, as they cover natural genetic variation in thousands of regulatory elements. For example, the SuRE data may be used to refine computational predictions of SNP effects ^12,28,40^.

Finally, it will be interesting to expand this type of analysis to individuals with a genetic disorder to capture additional disease-relevant variants that might not be found in the general population.

## Acknowledgements

We thank the NKI Genomics Core Facility for technical support and members of our laboratories for helpful discussions. Supported by ERC Advanced Grant (GoCADiSC) 694466 to B.v.S.; and ERC Starting Grant 637587 (HAP-PHEN) to E.d.W. The Oncode Institute is supported by KWF Dutch Cancer Society. HJB is supported by NIH grant R01HG003008 and Columbia University’s Vagelos Precision Medicine Pilot Program.

## Competing interest

J.v.A. declares a conflict of interest as the founder of Gen-X B.V. (www.gen-x.bio). E.d.W. is co-founder and shareholder of Cergentis.

## Author contributions

J.v.A. designed and performed experiments, analyzed data and wrote the manuscript. L.P., V.D.F. and H.J.B. developed algorithms and analyzed data. M.d.H. M.B., M.V., R.v.W., H.T., F.C., U.V., E.d.W. and L.F. generated and/or analyzed data. B.v.S. designed experiments, analyzed data and wrote the manuscript.

## ONLINE METHODS

### SuRE library preparation and barcode-to-fragment association

SuRE libraries were generated as described previously ^25^. DNA was isolated from lymphoblast cell lines HG02601, GM18983, HG01241 and HG03464 obtained from Coriell Institute, fragmented and gel-purified to obtain ~300bp elements. For each genome, two SuRE libraries were generated, each of an approximate complexity of 300 million fragment-barcode pairs. This was done by transformation in CloneCatcher DH5G electrocompetent Escherichia coli cells (#C810111; Genlantis) as done previously, or in E. cloni 10G (#60107-1; Lucigen). Barcode-to-fragment association was done as described previously ^25^, except that because of the smaller genomic insert size no digest with a frequent cutter was required. Thus, after I-CeuI digest and self-ligation we immediately proceeded to the I-SceI digest.

### Cell culture and transfection

K562 cells were cultured and transfected as described ^25^. HepG2 (#HB 8065; ATCC) were cultured according to supplier’s protocol and transiently transfected in the same manner as K562 cells except that program T-028 was used for nucleofection, 7.5 μg plasmid was used for each 5 million cells, and cells were harvested 48 hours after transfection. One hundred million cells were transfected for each replicate. Every 3 months all cells in culture were screened for mycoplasma using PCR (#6601; Takara).

### CRISPR-Cas9 mediated editing of rs3748136

We performed our CRISPR experiments on a K562 subclone in which *NR_125431* was active (subclone BL_2) because initial experiments revealed that in the K562 pool, *NR_125431* is expressed in only ~25% of the cells (Figure S4c). Five million of the BL_2 cells were nucleofected as described above with 2 μg of vector pX330-U6-Chimeric_BB-CBh-hSpCas9, a gift from Feng Zhang (Addgene plasmid # 42230 ^37^) encoding Cas9 and the chimeric guide RNA; and 20 pmol repair template (see **Supplemental Table 2** for nucleotide sequences of guide RNAs). Cells were then cultured for 3 days in the presence of 1μM DNA-PK inhibitor (#NU7441; Cayman Chemical Company) and then expanded for another 5 days without this inhibitor. For genotyping we used a PCR amplicon that that included SNP rs453301, ~250bp downstream of rs3748136 that was also heterozygous in K562. After confirming editing efficiency for the population of cells using Sanger sequencing and TIDE analysis ^41^, single cells were cloned out. After expansion, clones were genotyped on the single PCR amplicon, and classified as successfully edited when they were heterozygous (i.e. not edited) at rs453301 but homozygous for rs3748136, or they were classified as wild-type when both loci were still heterozygous. Of note, we identified many clones in which our CRISPR editing caused deletions around the targeted SNP; these were discarded. Successfully edited clones and wild-type clones were then analyzed for RNA expression by RT-qPCR. Since the chromosome that was edited at rs3748136 was the only chromosome showing expression of *NR_125431* to begin with, we are looking at the increased expression from that chromosome after editing (see main text), even though the RT-qPCR is not allele specific. See **Supplemental Table 2** for oligonucleotide sequences used.

### RT-qPCR

RNA was isolated from 1-5 million cells using Trisure (#BOI-38033; Bioline). DNase digestion was performed on ~ 1.5 μg RNA with 10 units DNase I for 30 min (#04716728001; Roche) and DNase I was inactivated by addition of 1 μl 25 mM EDTA and incubation at 70 °C for 10 min. cDNA was produced by adding 1 μl 50 ng/μl random hexamers and 1 μl dNTP (10mM each) and incubated for 5 minutes at 65°C. Then 4 μl of first strand buffer, 20 units RNase inhibitor (#EO0381; ThermoFisher Scientific), 1 μl of Tetro reverse transcriptase (#BIO-65050; Bioline) and 2 μl water was added and the reaction mix was incubated for 10 minutes at 25°C followed by 45 minutes at 45°C and heat-inactivation at 85° for 5 minutes. qPCR was performed on the Roche LightCycler480 II using the Sensifast SYBR No-ROX mix (#BIO-98020; Bioline). All expression levels were calculated using the 2^ΔΔCt^ method and normalized to the internal control GAPDH. See **Supplemental Table 2** for oligonucleotide sequences used for qPCR.

### Targeted Locus Analysis

TLA was performed essentially as described ^42^. Briefly, roughly 5 million K562 cells were cross-linked with 4% formaldehyde and cut with NlaIII. After ligation, the template was de-crosslinked and further digested with NspI. The second ligation yields circular DNA that is used as input for the inverse PCR reaction. We performed two PCR reactions: the first with primers adjacent to rs1053036, located in the last exon of the *NR_125431* and the second with primers adjacent to rs3748136, located in the intergenic region (**Supplemental Table 2**). The PCR amplicons were combined and we generated sequencing libraries using the KAPA High Throughput Library Preparation Kit (# 7961901001; Roche). We generated 2×150 sequences on an Illumina MiSeq. Sequence reads were mapped to hg19 using BWA-SW^43^. The resulting bam files and the K562 vcf file (obtained from whole genome sequencing at Novogene) were used as input for HapCUT2 ^44^ with the –hic option turned on to phase the variants.

### Cell culture and generation of nuclear extracts from K562 cells

Nuclear extracts were generated from K562 cells essentially as described ^45^. Briefly, cells were washed with PBS and then resuspended in 5 cell pellet volumes of hypotonic Buffer A (10 mM HEPES (pH 7.9), 1.5 mM MgCl2, 10 mM KCl). After incubation for 10 minutes at 4 °C, cells were collected by centrifugation and resuspended in two pellet volumes of buffer A supplemented with 0.15% NP40. Cells were then lysed by dounce homogenization using 35 strokes with a type B (tight) pestle on ice. Crude nuclei were collected by centrifugation and then lysed in two pellet volumes of Buffer C (420mMNaCl2, 20mMHEPES (pH 7.9), 20% (v/v) glycerol, 2 mM MgCl2, 0.2 mM EDTA, 0.1% NP40, EDTAfree complete protease inhibitors (Roche), and 0.5 mM DTT) by rotation for 1 h at 4 °C. After centrifugation for 20 minutes at 21.000 g, nuclear extract was collected as the soluble fraction. This extract was then aliquoted, snap-frozen and stored at -80 °C until further usage.

### DNA affinity purification and LC-MS analysis

Oligonucleotides for the DNA affinity purifications were ordered from Integrated DNA technologies with the forward strand containing a 5’ biotin moiety; see **Supplemental Table 2**). DNA affinity purifications and on-bead trypsin digestion was performed on 96-well filter plates essentially as described ^46^. Tryptic peptides from SNP variant pull-downs were desalted using Stage (stop and go extraction) tips and then subjected to stable isotope di-methyl labeling on the Stage tips ^34^. Matching light and heavy peptides were then combined and samples were finally subjected to LC-MS and subsequent data analyses using MaxQuant ^47^ and R essentially as described ^48^.

### External data sources

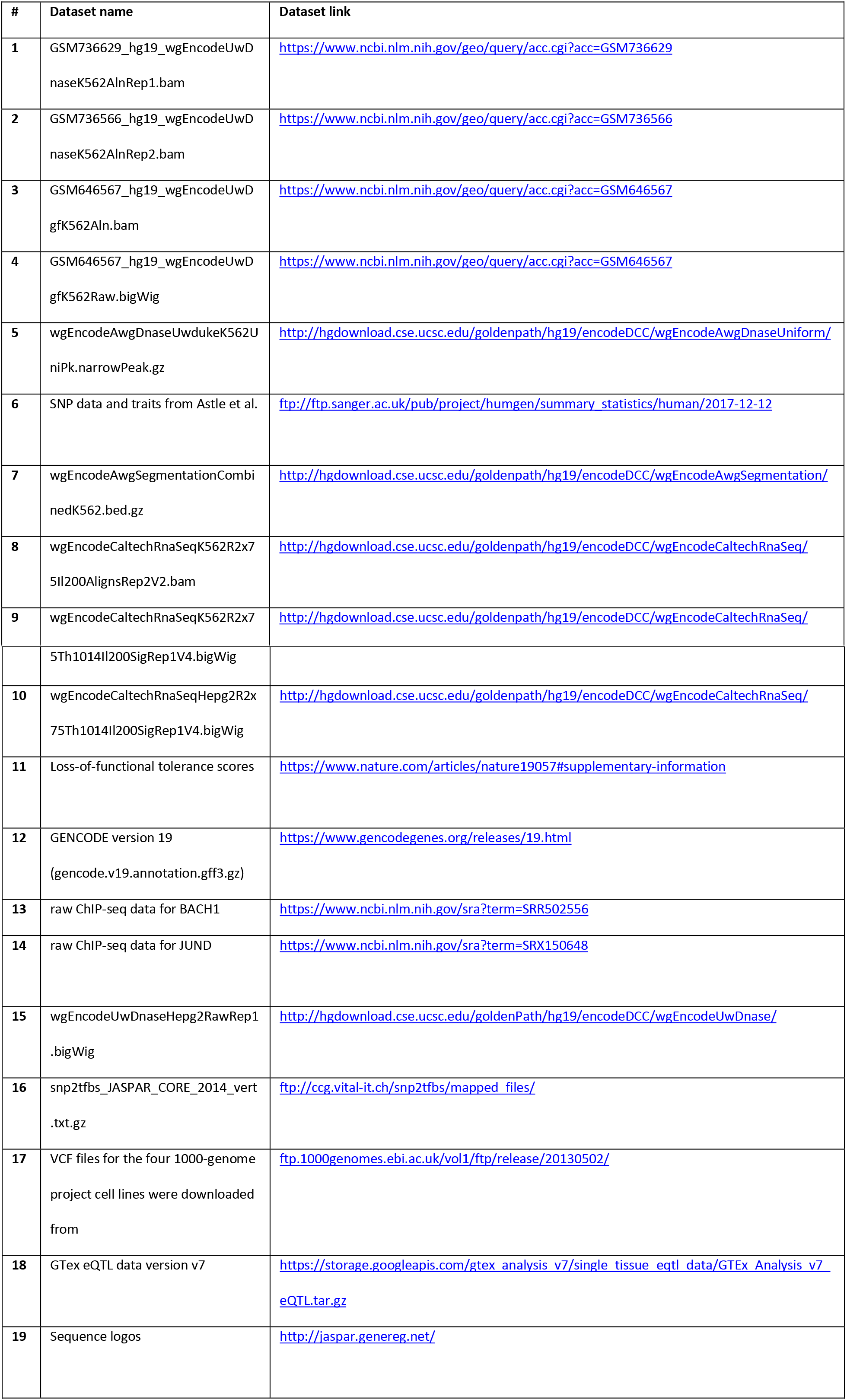

### Illumina sequencing

Paired end sequencing (150bp) of SuRE libraries was done by Novogene on the HiSeq-X platform, generating about 1 billion reads per library. Standard full genome sequencing and variant calling for the K562 cell line was done by Novogene on the HiSeq-X10 platform with PE 150bp reads amounting to approximately 100Gb or a ~30 fold coverage of the genome. Single end sequencing on reverse transcribed, PCR amplified barcodes was done by the NKI Genomics Core Facility on a HiSeq2500 machine.

### Sequencing data processing

Paired end reads (PE reads) of the SuRE libraries (for associating genomic positions and barcodes for each SuRE-fragment), and single end reads (SE reads) of the PCR amplified barcodes (representing raw SuRE expression data), were processed to remove adapter and vector backbone sequences, using *cutadapt* (V1.9.1, ^49^). All PE and SE reads were discarded if the barcode sequence contained Ns or the sequence was not exactly 20 nucleotides. The remaining sequences in the PE reads are combinations of barcode sequences and gDNA sequences, whereas the SE reads only yield barcode sequences. The latter barcodes are simply recorded and counted. The gDNA sequences of the SuRE libraries were mapped to the reference genome sequence (hg19, including only chr1-22, chrX), using *bowtie2* (V2.3.2, ^50^), with a maximum insert length set to 1kb. Read pairs with either the forward or the reverse gDNA sequence less than 6 nucleotides, and read pairs not aligned as ‘proper pair’, were discarded. To prevent allelic biases in alignment we used *WASP* ^51^ and SNP annotations from the 1000-genome project (external data source #17) to discard all reads potentially resulting in biased alignments.

The resulting associations of barcode sequence-genomic position pairs were further processed as follows:

1. Identical barcodes associated with multiple alignment positions were discarded except for the most abundant barcode-position pair.
2. Different barcodes associated with the exact same alignment position were merged; i.e. the barcode sequence associated with this position was set to the most frequent barcode sequence in the set, and the total number of PE reads in the set was used as count for this barcode-position pair.

Next, the barcodes identified in the SE reads were matched to the barcodes in the remaining barcode-position pairs, and *‘SuRE-count’* tables were generated associating barcode sequences, genomic positions, and counts for associated PE reads and matched SE reads for each of the biological replicates.

### SNP annotation

The fragments, specified in the SuRE-count tables were further annotated with SNP positions and base identities. For this annotation only SNPs were considered which were single nucleotide, bi-allelic over all 4 considered genomes. For each SNP in such a fragment we determined its base identity as observed in the actual sequence reads. As the region is defined by paired end reads some central part may not be covered by genomic sequence reads, in which case the uncovered SNPs were assigned the IUPAC representation of both allelic values. In the latter case, if the allelic value in the 2 parental chromosomes of the cell line were identical (based on annotation by the 1000 genome project, external data source #17), this inferred base identity was used for annotation.

### Generating BigWigs of SuRE enrichment profiles

For each strand separately, we determined the cDNA barcode count for all SuRE fragments overlapping a given position. This total was divided by the total counts of the SuRE fragments measured in SuRE library (iPCR barcode counts) to give the SuRE enrichment. Within each transfection replicate the genome-wide normalized SuRE enrichment was scaled to a mean of 1. Transfection replicates were then combined by summing the SuRE enrichment scores at each position, and scaling the resulting SuRE enrichment scores again to a genome-wide mean of 1. Then, the library replicates were also combined and again scaled to a genome-wide mean enrichment score of 1. Of these datasets BigWig files were generated. This analysis was done disregarding variants and is therefore independent from the identification of the raQTLs (see below).

### Re-mapping of BACH1 and JUND ChIP-seq data

Fastq files were downloaded from the SRA repository (external dataset #13, #14) using *fastq-dump* from the SRA-tools package. For BACH1 we downloaded data from data sets *SRR502556* and SRR502557, for JunD from data sets *SRR502542* and *SRR502543*. Reads were aligned to the human reference sequence (hg19, including only chr1-22, chrX), using bowtie2 (V2.3.2, ^50^) with default settings.

### Identification of raQTLs

First, for each transfection replicate the relative sequencing depth of cDNA barcodes was determined as the total cDNA barcode counts divided by the library complexity (i.e. the number of unique fragments identified in the library). The cDNA reads of all samples that were more than 1 standard deviation more deeply sequenced than the mean for all samples (i.e. all 3 K562 and HepG2 transfection replicates for library HG02601 library 1, and all HepG2 transfection replicates for HG02601 library 2) were downsampled to the mean relative cDNA read depth. This was done to avoid biases due to excessive differences in sequencing depth. Then, per chromosome, SuRE-count tables for each of the 8 SuRE libraries were expanded to contain for each fragment a normalized iPCR barcode count (normalized for iPCR sequencing depth by scaling the counts to reads per billion) and a normalized cDNA barcode count for the combined transfection replicates in K562 and HepG2 (normalized for cDNA sequencing depth by scaling the counts to reads per billion). Next a ratio of the normalized cDNA barcode count over normalized iPCR barcode count was calculated which we refer to as the SuRE signal (S_SuRE_) for that fragment. Since both values used to obtain this ratio are expressed as reads per billion, the resulting S_SuRE_ essentially represents an enrichment score.

Using this S_SuRE_ per fragment, a mean S_SuRE_ was calculated for each variant of each SNP as the mean S_SuRE_ for all fragments containing that fragment. Also, for each SNP a Wilcoxon test was performed comparing the S_SuRE_ of all fragments containing the REF allele with the S_SuRE_ of all fragments containing the ALT allele. The resulting vector of P-values (one for each SNP) is referred to as P_t_. In addition, to estimate the False Discovery Rate (FDR), the same Wilcoxon test was applied once after random shuffling of the S_SuRE_ of the fragments among the two variants, yielding a vector of P-values referred to as P_r_. We then focused on those SNPs for which both variants were covered by at least 10 fragments and no more than 999 fragments, and for which at least one of the variants had S_SuRE_ >4 (170,118 SNPs in K562; 395,756 SNPs in HepG2). For FDR=5%, we then chose the lowest P-value cutoff p_cut_ for which the number of SNPs with P_t_ < p_cut_ was at least 20 times larger than the number of SNPs with P_r_ < p_cut_. We refer to this set of SNPs as *raQTLs*. The same procedure was applied for data from K562 and HepG2 cells. For K562 p_cut_ was 0. 006192715 and for HepG2 p_cut_ was 0.00173121.

### Enrichment of raQTLs in ENCODE classes

For Figure 2a we used the GenomicRanges package of BioConductor ^52^ to determine the enrichment or depletion of raQTLs in ENCODE chromatin classes (external dataset #7) as compared to all 5.9 million SNPs we assessed.

### Comparison of SuRE to DNase-seq allelic imbalance

In Figure 2 we plotted the DNase-seq signal around the raQTLs using external data source #4 and BioConductor package CoverageView (version 1.4.0) and 25bp windows. Figure 2c was generated from the same data. For the analysis of allelic imbalance in the DNase-seq signal we combined three available experiments (external data source #1,#2,#3) and extracted from the bam-files the reads that overlapped a SNP that was found to be heterozygous in K562 in our own genome sequencing analysis (see ‘Illumina sequencing’ section). We focused on those raQTLs for which we found at least 20 DNase-seq reads and quantified the ratio of reads containing the REF allele over the reads containing the ALT allele, after adding for each allele a pseudocount of 1. Similarly, from our own genome sequencing of K562 we quantified the ratio of reads containing the REF allele over reads containing the ALT allele, after adding for each allele a pseudocount of 1. Finally, the DNase allelic imbalance was calculated as the DNase-seq allele-ratio over the genomic allele-ratio.

For the SuRE data, the allelic imbalance was calculated as the ratio of S_SuRE_ for the REF allele over the S_SuRE_ of the ALT allele (since both these values are already normalized for coverage in the libraries).

To obtain a matching set of control SNPs we intersected a DNase-peak annotation (external data source #7) with our SNPs, and we retrieved the DNase-seq variant counts for the 2500 overlapping SNPs with the highest SuRE P_t_ values. We required at least 20 reads covering the SNP, and from the resulting set we randomly selected a subset of SNPs of the same size as the set of raQTLs (Figure 2f). To this control set we applied the same analysis of allelic imbalance as to the raQTLs.

### Allele frequencies

MAFs of SNPs were obtained from the 1000 Genomes Project (external data source #17). Common variants were defined as SNPs with MAF > 0.05. Of the 5,919,293 SNPs in our SuRE dataset 4,569,323 classify as common variant. This is ~57% of the estimated 8 million common SNPs according to the 1000 Genomes Project ^1^. The proportion of raQTLs for which the SuRE effect could be resolved to a single SNP was calculated as the fraction of raQTLs for which neither neighbor SNP was also a raQTL.

### Motif disruptions

We made use of SNP2TFBS (external data source #16) ^28^ to identify all SNPs for which there was a difference in predicted affinity for a TF between the REF and ALT allele. For each SNP we only considered the TF listed first in SNP2TFBS (i.e., the TF with the biggest absolute difference in motif score). For the comparison of motif disruptions in K562 and HepG2 we identified all raQTLs of K562 and HepG2 that caused motif disruptions, we down-sampled the K562 disruptions to the same number as the HepG2 disruptions (since K562 has more raQTLs), and plotted the ratio of the counts (plus one pseudo-count) for the 7 most extreme ratios for each cell type. Sequence logos were obtained from external dataset #19.

### Compiling a set of control SNPs that are matched to the significant SNPs

For the analyses in figures S2 and 5a we compared the set of raQTLs to a control set of matching SNPs that was selected as follows. We ranked all SNPs, first: by the S_SuRE_ (rounded to whole numbers) of the strongest variant, second: by the number of fragments containing the least covered variant, and third: by the number of fragments containing the most covered variant. This is intended to rank them based on regulatory element activity (S_SuRE_) and our sensitivity to detect a significant difference (coverage of the variants). Then we identified the raQTLs along this ranking and selected both direct neighbors, removed the raQTLs and down-sampled the resulting set to yield the same number of matched SNPs as raQTLs.

### raQTL density around loss-of-function tolerant and loss-off-function intolerant genes

For the definition of LOF-tolerant and LOF-intolerant genes, we used external dataset #11 and the same cut-offs as previously reported ^31^, classifying genes as having pLI scores of >0.9 as intolerant and <0.1 as tolerant. We then matched these genes to GENCODE version 19 annotation (external data source #12) to identify the corresponding TSSs. Using the GenomicRanges package of BioConductor ^52^ we defined regions around these TSSs of +/- 100 kilobases and subtracted all possible exons (as defined in GENCODE version 19). In the resulting regions we then determined the fraction of all SNPs that is a raQTL, and the fraction that belonged to the set of matched control SNPs. This analysis was done for both LOF-intolerant and LOF-tolerant genes.

### Integration with eQTL data

GTEx eQTL data ^8^ (release v7; external data source #18) were downloaded on 27 January 2018. For whole blood we used the extracted file *Whole_Blood.v7.signif_variant_gene_pairs.txt.gz* and for liver we used *Liver.v7.signif_variant_gene_pairs.txt.gz*.

Gene annotation tracks in Figure 4 were generated by the Gviz package of BioConductor, function BiomartGeneRegionTrack(). ENCODE ^27^ DNase-seq data used in this figure are from external data source #4, #15.

### Integration with GWAS data

Overlap between SNPs identified in the GWAS study by Astle et al. ^35^, was obtained by searching for significant SuRE SNPs within 100 kb of each of the 6736 lead SNPs identifying 1238 lead SNPs within 100kb of at least one significant SuRE SNP (external data source #6). The window of 100 kb was chosen to be substantially larger than the typical size of an LD block. For these lead SNPs we calculated the distance to SNP with the lowest P-value in our SuRE data (Figure 5a). As a control we did the same procedure for the set of matched SuRE SNPs. In Figure 5 only those SNPs with significant P-values in the GWAS study (cutoff: p < 8.31E-9 ^35^) are shown. Note that what we refer to as ‘lead SNPs’ are called ‘conditionally independent index-variant associations’ in the original GWAS study ^35^. Gene annotation tracks in Figure 5 were generated by the Gviz package of BioConductor, function BiomartGeneRegionTrack(). DNase-seq data used are from external data source #4.

### General data analysis and visualization

Data analysis and figure production was mostly done using various R (https://www.R-project.org) and BioConductor packages. Scripts are available upon request.

## Data availability

Raw sequencing data will be made available at GEO (currently uploading; ETA ~20^th^ of November). SuRE count tables, BigWig files for visualization of SuRE data tracks in genome browsers, and a table with SuRE data for all 5.9 million SNPs is available from OSF:https://doi.org/10.17605/OSF.IO/W5BZQ

## SUPPLEMENTAL DATA

– Supplemental Table 1: overview complexities and sequencing depth for all SuRE libraries

– Supplemental Table 2: oligonucleotide sequences

**Figure S1.**
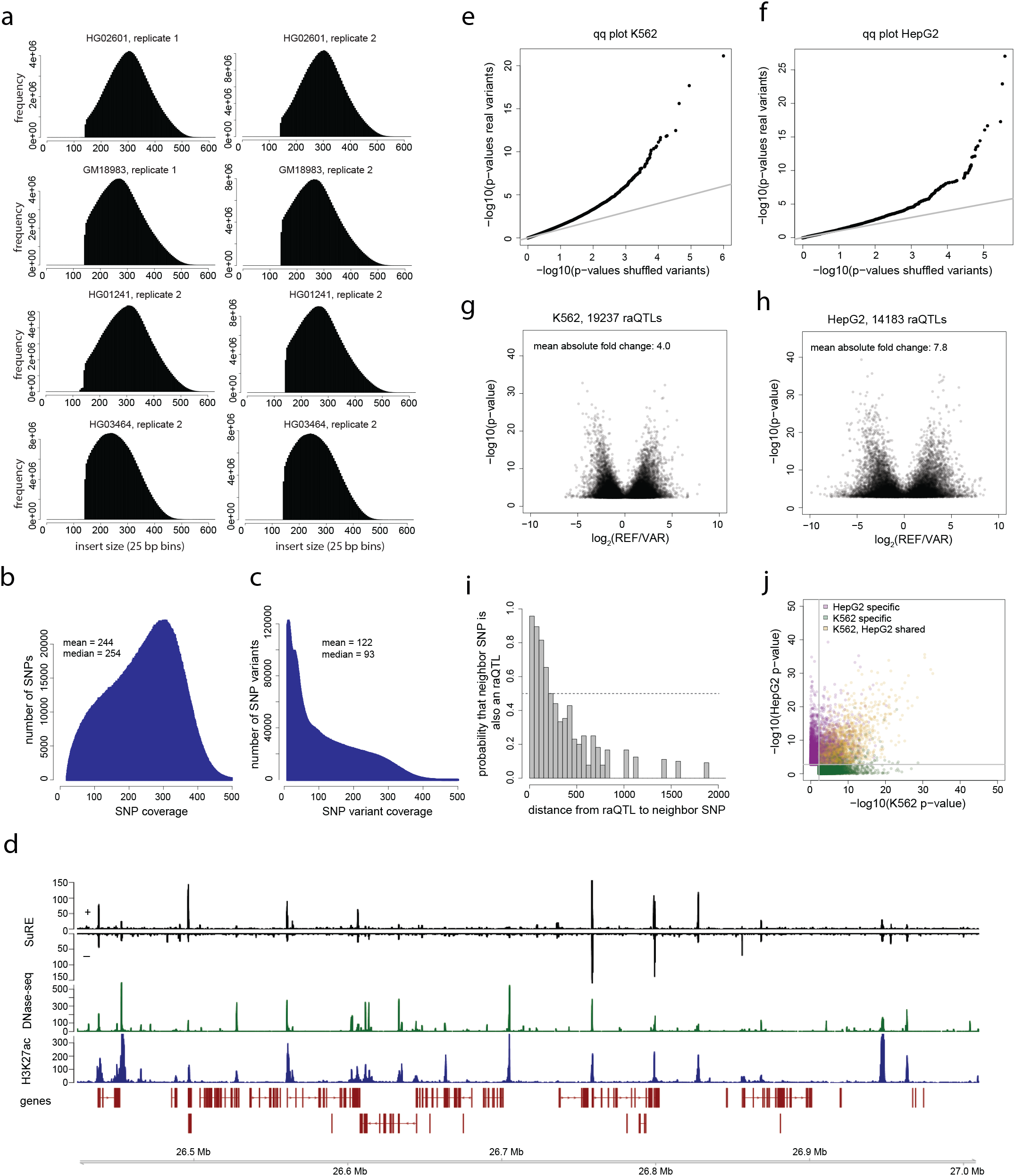
Characterization of SuRE libraries and SuRE data. **a**. Inserted fragment size distribution for each SuRE library (bin size 25 bp). **b**. Histogram showing the coverage of each SNP position in the combined SuRE libraries. **c**. Same as (**b**) but now for each SNP variant. **d**. Representative ~0.5-Mb genomic region showing SuRE signals of HG02601, SuRE library 1 in K562 cells, together with DNase-seq and H3K27ac signals ^27^ in K562 cells. **e**. qq plot showing the distribution of P-values for SNPs in SuRE in K562 (y-axis) compared to distribution of P-values obtained after random shuffling the SuRE expression values for each SNP (x-axis). Shown is a random subset of 100,000 SNPs. Gray line indicates y=x diagonal. **f**. Same as (**e**) but for HepG2. **g**. Volcano plot showing for all raQTLs in K562 the log_2_ difference in SuRE signals for the REF and the ALT allele (x-axis) and the associated P-values (y-axis). **h**. Same as (**g**) but for HepG2. **i**. Histogram showing for all raQTLs in K562 the probability of the nearest neighbor SNP also being a raQTL, as a function of their distance. The dotted gray line indicates probability 0.5. **j**. SuRE P-values in K562 and HepG2 cells for all SNPs that are raQTLs in at least one of the two cell types. Gray lines indicate the P-value cut-offs for each cell type.

**Figure S2.**
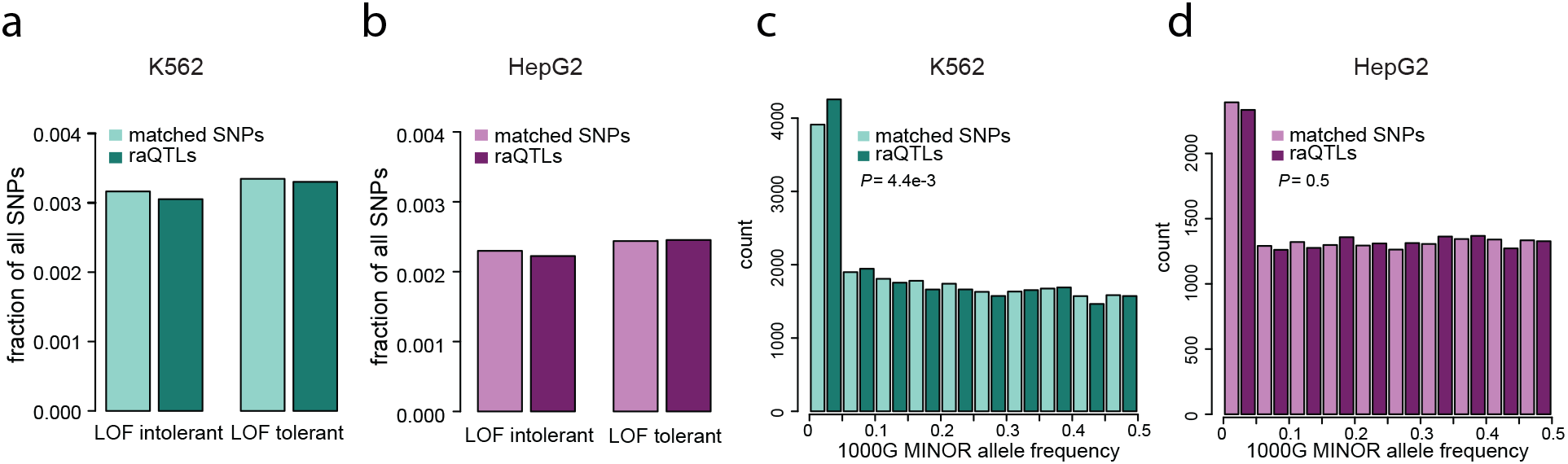
Genomic distributions and minor allele frequencies of raQTLs. **a**. Frequencies of raQTLs in K562 cells (dark color) or matching control SNPs (pale color) among all non-exonic SNPs within 100 kb of TSSs of loss-of-function tolerant genes or loss-of-function intolerant genes^31^. **b**. Same as (**a**) but for HepG2 cells. **c**. Distributions of minor allele frequencies according to the 1000 Genomes Project ^1^ for raQTLs (dark color) and matched control SNPs (pale color) in K562 cells. **d**. Same as (**c**) but for HepG2.

**Figure S3.**
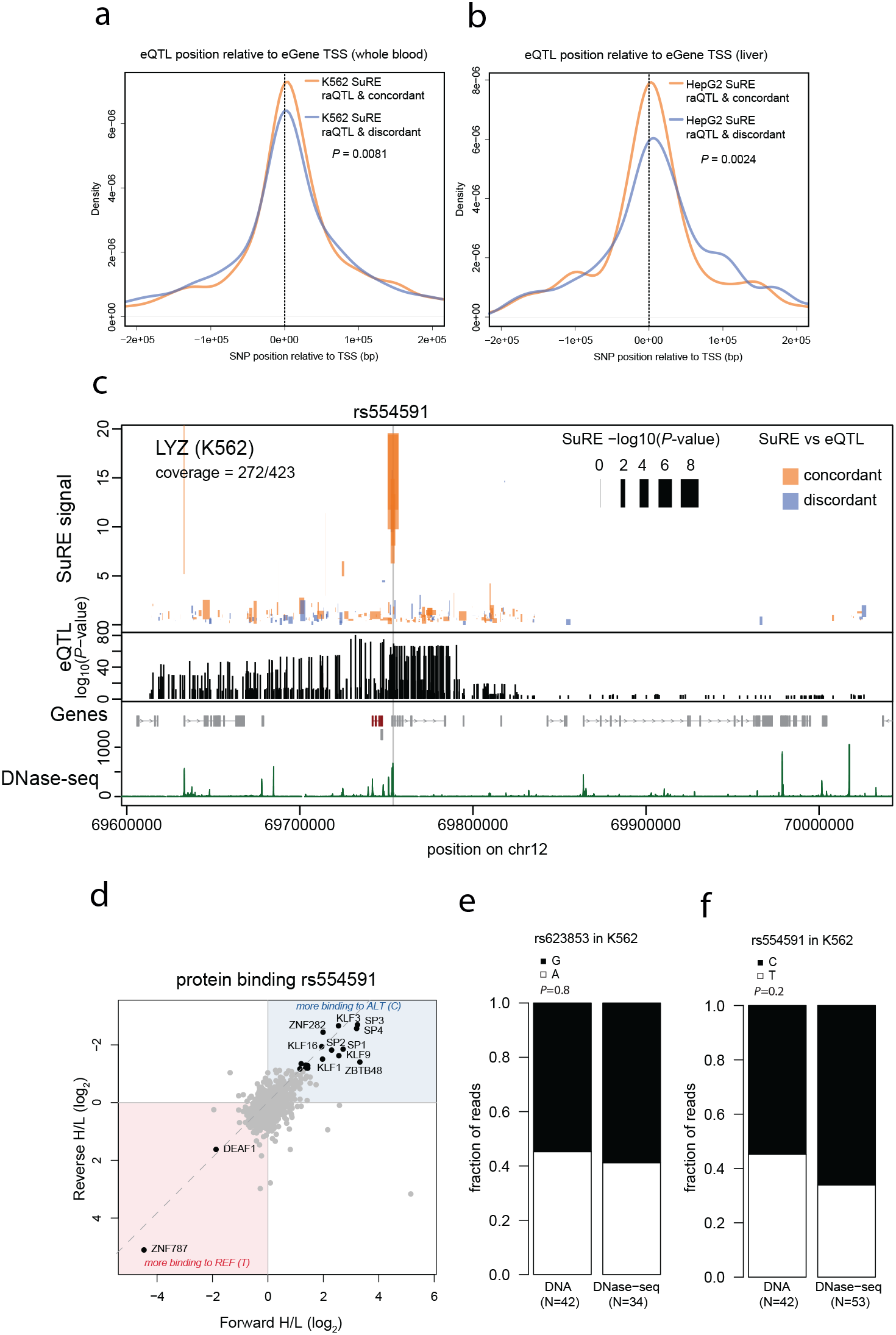
Additional data related to eQTL – SuRE comparisons. **a**. Distributions of distances between whole blood eQTLs and the TSS of the eGenes for concordant (orange) and discordant (blue) raQTLs in K562. **b**. Same as (**a**) but for liver eQTLs and SNPs that are raQTLs in HepG2. **c**. Genome track plot combining SuRE data and eQTL mapping data for *LYZ* in whole blood, similar to main Figure 4e. **d**. Protein binding analysis for rs554591, similar to main Figure 4f. **e**. Barplots indicating fraction of reads containing each of the two variants for rs623853 in K562 genomic DNA (left) and K562 DNase-seq reads ^27^ (right). **f**. Same as (**e**) but for rs554591.

**Figure S4.**
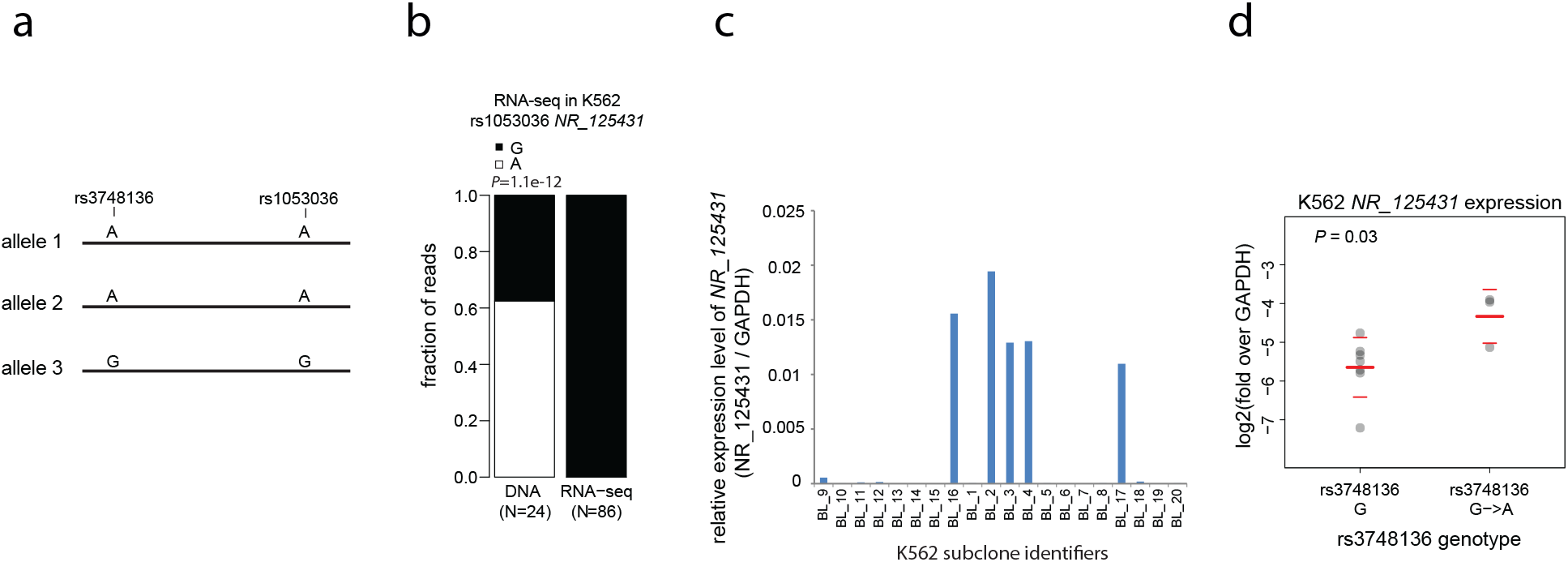
Unexplained allele-specific variation of *NR_125431* expression before and after editing of rs3748136. **a**. The A and G alleles of rs1053036 in *NR_125431* are cis-linked to the A and G alleles of rs3748136, respectively. Linkage model is based on TLA mapping. This locus in K562 cells is most likely triploid. **b**. Fraction of reads containing each of the two variants of SNP rs1053036 in *NR_125431* in K562 genomic DNA (left) and K562 RNA-seq reads (right). The complete lack of expression of the A allele of *NR_125431* is unexpected and may point to a genetic defect of the A allele in K562 cells. **c**. Clonal lines derived from K562 cells show extreme expression variation of *NR_125431*. For CRISPR-based editing we proceeded with clone BL_2. **d**. Expression of *NR_125431* in subclones derived from clone BL_2 subjected to CRISPR/Cas9 editing of rs1053036. Seven unaltered subclones and three G->A edited subclones were assayed by RT-qPCR of *NR_125431* (normalized to *GAPDH)*. Red lines indicate means and thin red lines indicate standard deviations.

